# A novel degenerate primer set for eDNA metabarcoding of amphibians, turtles and fish

**DOI:** 10.1101/2024.09.27.615469

**Authors:** Torrey W. Rodgers, John R. Olson, Charles P Hawkins, Karen E. Mock

## Abstract

Metabarcoding of environmental DNA (eDNA) has revolutionized the detection of aquatic species across large geographic scales. However, the effectiveness of eDNA metabarcoding across taxa is marker-dependent, often requiring multiple markers to detect divergent taxonomic groups. Here, we introduce a novel metabarcoding marker designed to simultaneously detect amphibians, turtles, and fish from eDNA samples. We initially optimized this marker to match species from central and southern California, USA, and conducted validation on 525 field-collected eDNA samples from central California, while also testing an existing published amphibian marker from the literature. Additionally, we assessed the global applicability of the marker with *in-silico* analysis through comparison with 11,350 amphibian, turtle, and fish mitochondrial genomes downloaded from GenBank. Field validation demonstrated the markers robustness in detecting aquatic vertebrates, outperforming the published marker in sensitivity, particularly for turtles and fish. In-silico evaluation demonstrated the potential for global use of this marker, indicating broad applicability for amplifying eDNA from amphibians, turtles, and fish worldwide. Our findings highlight the utility of this novel marker for monitoring of aquatic vertebrate communities, with implications for conservation, management, and ecological research.

## Introduction

Metabarcoding of environmental DNA (eDNA) has revolutionized our ability to detect aquatic species from waterbodies across large geographic scales, and in many cases, eDNA sampling results in higher detection probability than traditional methods (Fediajevaite et al., 2021). One of the principal benefits of metabarcoding is that many species can be detected simultaneously from the same sample. Which species are detected, and which are missed, however, is highly dependent on the metabarcoding marker used, as well as the availability of a reliable reference sequence database for the marker of choice. Unfortunately, there is no perfect marker that can detect all species simultaneously, and as a result, it is often necessary to run multiple markers to detect multiple groups of organisms. For the sake of efficiency, however, metabarcoding markers that are broad enough to detect multiple groups of organisms simultaneously are highly advantageous. Several general markers exist for detection of a wide range of metazoan taxa including invertebrates (Geller et al., 2013; Leray et al., 2013; Rennstam Rubbmark et al., 2018; Tournayre et al., 2024); however, because they are so broad, these markers also amplify DNA form non-target taxa such as fungi, algae, and bacteria, such that sequence data representing vertebrate taxa often represents just a tiny fraction of total data, making these markers inefficient, and likely to miss rare vertebrate species, where the primary goal is detection of aquatic vertebrates.

To this end, we designed a novel metabarcoding marker to detect amphibians, turtles, and fish from eDNA samples simultaneously. Initially, we designed this marker to perfectly match all amphibian and turtle species from southern and central California, USA; however, we also demonstrate that this marker should perform well in amplifying DNA from most amphibian, turtle, and fish species worldwide. We tested the marker on pools of amphibian and turtle tissue samples, and on field-collected eDNA samples from lakes, rivers, wetlands, and streams in coastal central California. Additionally, we ran an existing published marker designed to detect anurans and salamanders (Valentini et al., 2016) on our field collected eDNA samples, to determine if our novel marker could detect amphibians as effectively as the existing marker while also detecting turtles and fish. Finally, we performed an *in-silico* analysis using downloaded mitochondrial genomes of amphibians, turtles, and fish from around the world to determine the applicability of our maker for use globally.

## Methods

### Marker Development

Reference sequence data from the mitochondrial gene 16S for all amphibians and turtle species present in coastal central and southern California was obtained from NCBI GenBank or generated via Sanger sequencing of tissue samples (table S1). Based on these sequence data, we designed a primer set optimized for detecting all amphibian and turtle species present in central and southern California. To design this marker, we modified an existing universal primer set (L2513 and H2714; Kitano et al., 2007) that has been previously used for eDNA detection of amphibians and fish (Evans et al., 2016). Primers L2513 and H2714 possessed numerous mismatches with our target species, particularly turtle species, including near the 3’ end of primer L2513 that would impede amplification or reduce sensitivity for detecting our target species. Thus, we initially introduced 3 degeneracies into the existing primers (this modified primer set was named marker 16S-mod-turtle, Table 1), one at the 20^th^ position of primer L2513 (re-named L2513-mod-turtle; Table 1), and two at the 6^th^ and 7^th^ position of primer H2714 (re-named H2714-mod-turtle; Table 1). After tissue DNA from California tiger salamander (*Ambystoma californiense)* failed to amplify in our tissue DNA primer testing phase (see below), we added an additional degeneracy to primer L2513-mod-turtle at position 21 to perfectly match available *A. californiense* sequences (re-named L2513-mod-tuTS; Table 1), and this primer was used in our marker (16S-AmTu; Table 1) for eDNA sample analysis.

**Table 1.**
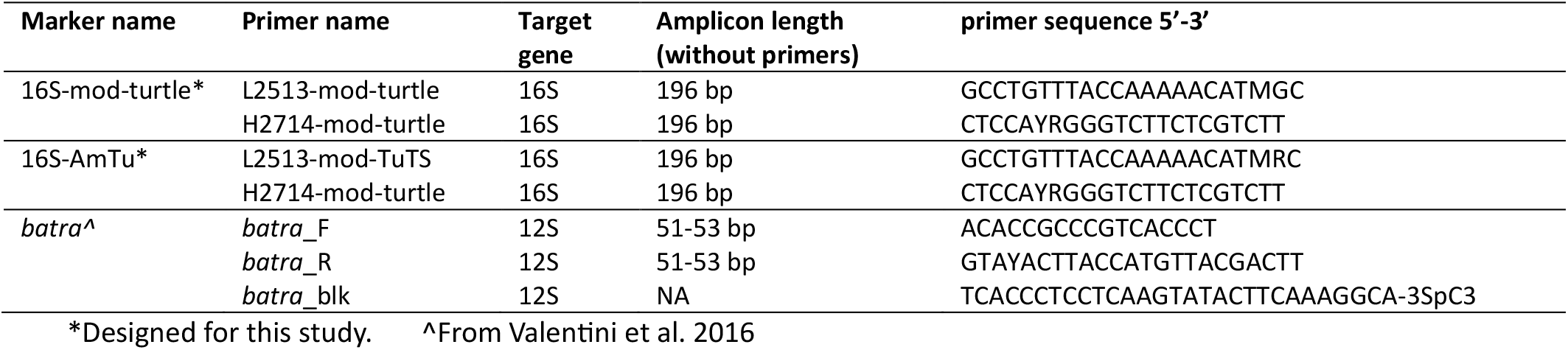
Primers used for metabarcoding of amphibians and turtles from eDNA samples.

### Tissue sample testing

We tested primer set 16S-mod-turtle (Table 1) with metabarcoding on a pool of tissue DNA samples from 20 California amphibian and turtle species from our study area (Table S2). PCR reactions contained 12.5 μl Amplitaq Gold 360 Master Mix (Thermo-Fisher Scientific) 0.2 μM of forward and reverse primer containing Illumina sequencing adapters, 0.2 μg/μl of bovine serum albumin, and 2 μl of tissue pool template in a total volume of 25 μl. Cycling conditions were 95° C for 10 minutes followed by 45 cycles of 95° C for 15 seconds, 55° C for 30 seconds, and 72° C for 30 seconds, followed by a final extension at 72° for 7 minutes. PCR products were then diluted 25:1 with sterile H_2_O, and this diluted PCR product was used to seed an indexing PCR utilizing Unique Dual Index (UDI) adapters for sample identification (Integrated DNA technologies). Indexing reactions contained 10 μl Amplitaq Gold 360 Master Mix (Thermo-Fisher Scientific), 1 μl of forward and reverse primer mix with UDI adapters, and 2 μl of pooled and diluted PCR product from the initial PCR in a total reaction volume of 20 μl. Cycling conditions for the indexing PCR were 95° C for 10 minutes followed by 15 cycles of 95° C for 15 seconds, 50° C for 30 seconds, and 72° C for 30 seconds, followed by a final extension at 72° for 10 minutes. PCR products were cleaned up and normalized with a SequalPrep™ Normalization Plate Kit (Thermo Fisher Scientific) and pooled for sequencing. The library was sequenced on Illumina MiSeq with the v2-500 cycle Nano Reagent Kit.

Additionally, we tested primer set 16S-AmTu (Table 1) on a pool of ‘exotic’ tissue DNA from six species including three anurans, two turtles, and one newt, from Africa or Asia (Table S3). This tissue pool was used as a positive control during sequencing of eDNA samples and was sequenced as described below.

### eDNA sample collection

Environmental DNA samples were collected following the protocol outlined in Carim et al. (2016) with modifications described below. We collected triplicate eDNA samples from 175 sites at 132 separate waterbodies (lakes were sampled at multiple sites) from March 29 – May 5, 2019, on US Department of Defense (DOD) and Los Padres National Forest (LPNF) lands across the California central coast (Fig. 1) Sites were distributed evenly across ownership, waterbody type, and environmental gradients such as climate, vegetation and fire history. We additionally collected 12 field blanks to monitor for cross contamination. Of the 175 sites sampled, 106 (61%) were on DOD land and 69 (39%) were within the LPNF. Of the 106 sites sampled on DOD lands, 31 sites were sampled on Vandenberg Air Force Base, 61 on Fort Hunter-Liggett, 6 on Camp Roberts, and 8 on Camp San Luis Obispo. Sites included 54 streams (31% of sites), 41 wetlands (23% of sites), and 80 sites distributed across 37 ponds/lakes. Sites were well distributed spatially and environmentally, but low-precipitation, low-elevation (i.e., coastal) sites were under sampled compared to the other environments because many of these systems were too dry to sample when field work was conducted.

**Figure 1.**
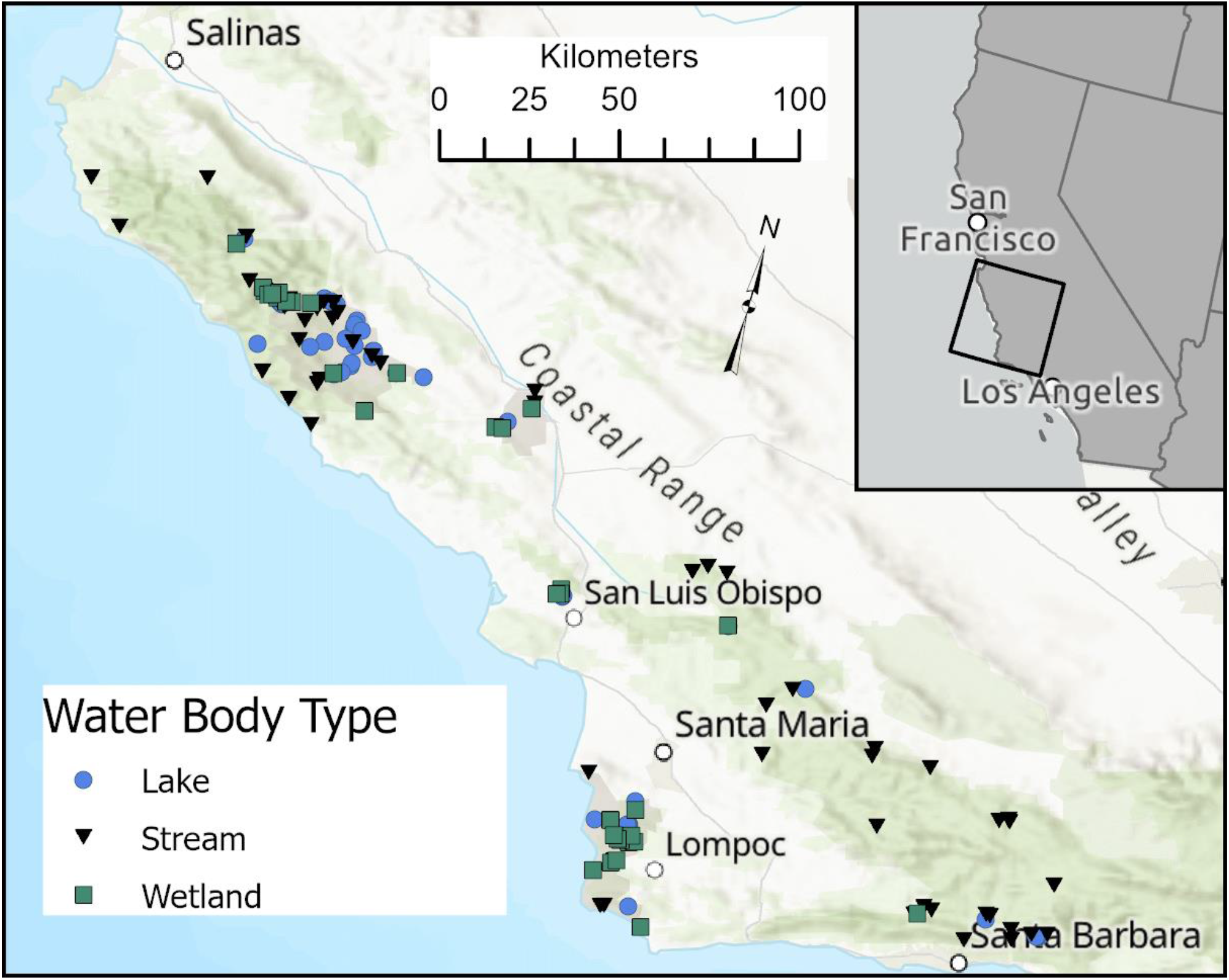
Locations of environmental DNA samples collected for field validation of metabarcoding markers.

At each sample location, crews pumped 1-5 liters of water through a Whatman 1827-047 1.5 μm glass microfiber filter with a battery-powered Geotech Geopump™ Peristaltic DC Pump (Geotech Environmental; Denver, CO). Lake and wetland samples were collected within 3 m of shore by placing the filter just below surface to avoid stirring up bottom sediments. Stream samples were collected by placing the filter intake on or near the stream bottom. Pumping continued until 5 L of water was filtered or until clogging. If pumping slowed to > 15 seconds between drops, the filter was considered to be clogged. If < 1 L was sampled prior to clogging, then a second filter was collected, and both filters were extracted together. Individual filters were carefully removed in the field with fresh sterile forceps and preserved in a 50-mL Falcon tube with silica gel desiccant. All sampling equipment was bleach-sterilized between each waterbody to eliminate cross-site contamination. Field blanks were collected after every 10th site (n = 12) by applying the collection protocol to one L of distilled water brought from the lab, as negative controls for potential field-based contamination. Samples were initially stored at room temperature for 8 to 13 weeks until all samples were collected and then shipped to the Molecular Ecology Laboratory at Utah State University where they were stored in a -20 freezer until extraction.

We collected 3 samples from each river or wetland site, but we sampled 1-3 sites at each lake depending on lake size to improve detections in large, unmixed waterbodies. We sampled lakes <20 m across at 1 site (i.e., 3 samples) lakes 20-30 m across at 2 sites (6 samples) and lakes > 30 m across at 3 sites (9 samples). When 2 or 3 sites were sampled, the additional sites were placed equidistantly around the lake. In total Environmental DNA samples were collected in triplicate from 175 sites at 132 separate waterbodies resulting in a total of 525 field-collected eDNA samples.

### eDNA Sample analysis

Samples were extracted in a dedicated room with the DNeasy Blood & Tissue Kit (Qiagen, Inc. Valencia, CA, USA) following a modified protocol (Rodgers et al. 2023). Each round of eDNA extraction (n=39) included one extraction blank negative control consisting of a clean, unused filter. All eDNA samples were amplified for next generation Illumina amplicon sequencing (metabarcoding) with two markers, our novel primer set 16S-AmTu (Table 1) and a published marker targeting the mitochondrial gene 12S (*batra;* Valentini et al. 2016; Table 1). For both markers, all samples were run in triplicate PCR. Reactions contained 2X (12.5 μl) Amplitaq Gold 360 Master Mix (Thermo-Fisher Scientific), 0.2 μM of forward and reverse primer containing illumina sequencing adapters, 0.2 μg/μl of bovine serum albumin, 4 μM of human blocking primer (*batra* reactions only), and 4 μl of eDNA template in a total volume of 25 μl. Cycling conditions were 95° C for 10 minutes followed by 45 cycles of 95° C for 15 seconds, 55° C (16S-AmTu) or 60° C (*batra*) for 30 seconds, and 72° C for 30 seconds, followed by a final extension at 72° for 7 minutes. After the initial marker-specific PCRs, triplicate PCR products were pooled by sample and diluted 25:1 with sterile H_2_O. This diluted PCR product was used to seed an indexing PCR utilizing Unique Dual Index (UDI) adapters for sample identification (Integrated DNA technologies). Indexing reactions contained 2X (10 μl) Amplitaq Gold 360 Master Mix (Thermo-Fisher Scientific), 1 μl of UDI primer, and 2 μl of pooled and diluted PCR product from the initial PCR in a total reaction volume of 20 μl. Cycling conditions for the indexing PCR were 95° C for 10 minutes followed by 15 cycles of 95° C for 15 seconds, 50° C for 30 seconds, and 72° C for 30 seconds, followed by a final extension at 72° for 10 minutes. PCR products were then cleaned-up and normalized with a SequalPrep™ Normalization Plate Kit (Thermo Fisher Scientific) and pooled for sequencing. Three separate libraries were created for each marker, each containing one field-replicate sample from each site. Each library contained a total of 192 samples, composed of 186 eDNA samples (including field blank and extraction negative controls), two positive control samples composed of a mix of tissue DNA from five exotic amphibian and turtle species not native North America, and four no-template negative control reactions. Each library (n=6) was sequenced on Illumina MiSeq with the v3-600 cycle (25 million read yield) reagent kit.

Bioinformatic processing of MiSeq data proceeded as follows. After demultiplexing, primer sequences were removed with CUTADAPT v. 1.18 (Martin, 2011). Next, data were filtered and denoised, pairedends were merged, and chimeras were removed with DADA2 (Callahan et al., 2016) within the QIIME2 environment (Bolyen et al., 2019). We then used ncbi BLAST (Altschul et al., 1990) to separate ASVs (Actual Sequence Variants) of aquatic amphibians, turtles, and fish from those of non-target taxa (e.g., ASVs from bacteria, fungus, mammals, and birds). For the ultimate taxonomic assignment of amphibian, turtle, and fish ASVs, we used the Naive Bayes classifier Scikit-learn (Pedregosa et al., 2011) within the QIIME2 environment against a custom reference sequence database containing sequences from all amphibians, turtles, and fish species known from our study area generated from reference tissue sequencing and additional sequences obtained from NCBI GenBank.

### In-silico primer evaluation

In addition to testing our novel marker on tissue and eDNA samples from California, we performed *in-silico* testing of primers from our 16S-AmTu marker to evaluate how well the marker is likely to perform for amplification of DNA from amphibian, turtle, and fish species globally. For this aim, we used the *in-silico* primer development and validation R package PrimerMiner (Elbrecht & Leese, 2017). We used PrimerMiner to download all complete amphibian and turtle mitochondrial genomes from NCBI GenBank. For fish, we downloaded mitochondrial genomes with PrimerMiner from the 8 most species rich fish orders (*Cypriniformes, Siluriformes, Characiformes, Cichliformes, Cyprinodontiformes, Gobiiformes, Perciformes*, and *Anabantiformes*), as these eight orders contain nearly 90% of all fish species (IUCN/SSC Freshwater Fish Specialist Group, 2024). Additionally, we included the order *Salmoniformes*, as this order is often of economic importance. We chose to use only complete mitochondrial genomes because the priming site of primer L2513-mod-TuTS is a conserved priming site that is very commonly used for sequencing of the 16S gene for amphibians and turtles. Thus, the majority of partial 16S sequences in Genbank do not include this priming region. Next, we used PrimerMiner to extract all complete 16S sequences from the mitochondrial genomes, and we clustered 16S sequences into Operational Taxonomic Units (OTUs) at 97% clustering to retain sequences representative of individual species. We then aligned the resulting 16S sequences using Sequencher software (version 5.4.6) and used the plot function in PrimerMiner to plot our 16S alignment for visual comparison with our primer sequences (Elbrecht & Leese, 2017).

## Results

### Tissue sample testing

Metabarcoding of DNA tissue pools of California amphibian and turtle species with primer set 16S-mod-turtle retrieved sequences from 18/20 species whose DNA was included in the pools. The two species not detected were *Pseudacris sierra and Ambystoma californiense*. The non-amplification of *P. sierra* was puzzling, as the primers match perfectly with available *P. sierra* sequences. However, *P. sierra* was the most commonly detected species from our eDNA samples (Table 2) and thus our primers are clearly capable of detecting this species. Upon closer examination of available *A. californiense* sequences, we noticed a mismatch with primer L2513-mod-turtle 2 bp from the 3’ end, a position which is highly likely to impede amplification. Thus, we further modified primer L2513-mod-turtle with an additional degeneracy at this position (renamed L2513-mod-tuTS; Table 1) to perfectly match available *A. californiense* sequences, and this new primer set (16S-AmTu, Table 1) was used for eDNA analysis. Additionally, primer set 16S-AmTu amplified DNA from all six ‘exotic’ species in our mixed positive control sample, which included tissue DNA from three anuran species, two turtle species, and one newt species, all from Africa or Asia (Table S3).

**Table 2.** Amphibian and turtle species detected by two metabarcoding markers from environmental DNA samples collected from central California. 16S-AmTu is s marker designed for this study, and *batra* is a marker designed by Valentini et al., (2016).

| Species | Common name | # Samples |  |
| --- | --- | --- | --- |
|  |  | 16S-AmTu | *batra* |
| *Pseudacris hypochondriaca/sierra* | Baja California/sierran chorus frog | 247 | 193 |
| *Anaxyrus boreas* | Western toad | 91 | 116 |
| *Lithobates catesbeianus* | American bullfrog | 69 | 35 |
| *Actinemys marmorata* | Western pond turtle | 47 | 5 |
| *Taricha torosa* | California newt | 27 | 16 |
| *Rana draytonii* | California red-legged frog | 26 | 33 |
| *Spea hammondi* | Western spadefoot | 26 | 5 |
| *Ambystoma californiense/tigrinum* | Tiger salamander/CA tiger salamander | 20 | 18 |
| *Trachemys scripta* | Pond slider turtle | 10 | 0 |
| *Pseudacris cadaverine* | California tree frog | 8 | 11 |
| *Xenopus laevis* | African clawed frog | 6 | 0 |
| *Rana boylei* | foothill yellow-legged frog | 5 | 2 |
| *Anaxyrus californicus* | Arroyo toad | 3 | 0 |
| *Chrysemys picta* | Painted turtle | 1 | 1 |

### eDNA Sample analysis

Our metabarcoding analysis of field-collected eDNA samples detected 14 amphibian and turtle species. These species included three turtles, nine anurans, one newt, and one salamander (Table 2). All 14 of these species were detected by our novel 16S-AmTu marker. The published *batra* marker from Valentini et al. (2016) detected 11 total amphibians and turtle species, including seven anurans, two turtles, one newt, and one salamander. Our novel marker detected most species from a greater number of samples than the published *batra* marker (Table 2), except in the case of California Red-legged Frog *Rana draytonii* (33 samples with *batra* vs. 26 samples with 16S-AmTu), western toad *Anaxyrus boreas* (116 samples with *batra* vs. 91 samples with 16S-AmTu), and California tree frog *Pseudacris cadaverina* (11 samples with batra vs. 8 samples with 16S-AmTu). Discrepancy in rates of detection was most notable for turtle species. Native western pond turtle *Actinemys marmorata* was detected in 47 samples with 16S-AmTu vs. just five for *batra*, and the invasive pond slider *Trachemys scripta* was detected in 10 samples with 16S-AmTu and was not detected at all with the *batra* marker. A discrepancy was also seen between the two markers for two threatened and endangered anuran species. Western spadefoot toad was detected in 26 samples with 16S-AmTu vs. just five for the *batra* marker, and arroyo toad *Anaxyrus californicus* was detected in three samples with 16S-AmTu, and not detected at all with the *batra* marker. Neither marker could distinguish between Baja California chorus frog *Pseudacris hypochondriaca* and Sierran chorus frog *Pseudacris sierra*; however, this is not surprising, as these species are very closely related, and were recently split from the Pacific chorus frog *Pseudacris regilla* (Recuero et al., 2006). Likewise, neither marker was able to distinguish between the closely related California tiger salamander *Ambystoma californiense* and eastern tiger salamander *Ambystoma tigrinum*. This latter result is important because the California tiger salamander is a vulnerable species that is threatened by hybridization with the invasive eastern tiger salamander (Ryan et al., 2009). Mitochondrial markers also cannot distinguish hybrids from parent species because of maternal inheritance of mitochondrial DNA.

Our 16S-AmTu marker detected a total of 21 fish species from 9 orders and 12 families (Table 3). The *batra* marker also amplified DNA from fish, but taxonomic power was low for assigning fish taxa from sequences for *batra*, and thus we only attempted taxonomic assignment for fish from the 16S-AmTu marker. Additionally, the 16S-AmTu marker incidentally detected 27 mammal species from our eDNA samples (Table S4) as well as some reptile and bird taxa.

**Table 3.** Fish species detected by the metabarcoding marker 16S-AmTu from environmental DNA samples collected from central California.

| **Species** | **Common Name** | **# Samples** |
| --- | --- | --- |
| *Dorosoma petenense* | Threadfin shad | 7 |
| *Catostomus sp.* | Sucker Sp. | 57 |
| *Cyprinus carpio* | Common carp | 32 |
| *Lavinia exilicauda* | Hitch | 50 |
| *Pimephales promelas* | Fathead minnow | 4 |
| *Gila orcutti* | Arroyo chub | 1 |
| *Rhinichthys osculus* | Speckled dace | 25 |
| *Gambusia affinis* | Western mosquitofish | 24 |
| *Gasterosteus aculeatus* | Three-spined stickleback | 26 |
| *Eucyclogobius newberryi* | Northern tidewater goby | 4 |
| *Lepomis cyanellus* | Green sunfish | 16 |
| *Lepomis macrochirus* | Bluegill | 29 |
| *Micropterus dolomieu* | Smallmouth bass | 8 |
| *Micropterus salmoides* | Largemouth bass | 41 |
| *Lampetra tridentata* | Pacific lamprey | 12 |
| *Oncorhynchus mykiss* | Rainbow trout | 42 |
| *Salmo trutta* | Brown Trout | 1 |
| *Cottus asper* | prickly sculpin | 30 |
| *Ameiurus sp.* | Bullhead sp. | 4 |
| *Ictalurus punctatus* | Channel catfish | 6 |

### In-silico primer evaluation

We used PrimerMiner (Elbrecht & Leese, 2017) to download a total of 2767 complete amphibian and turtle mitochondrial genomes, 2189 from amphibians (1511 from the Family Anura, 595 from the Family Caudata, and 83 from the Family Gymnophiona) and 219 from turtles. These were clustered into 1472 OTUs representative of individual species (Figure 2). Visual inspection of Figure 2 demonstrates that both primer sets match nearly all amphibian and turtle reference sequences perfectly, with the exception of some mismatches of Testudines and Gymnophiona sequences at the far 5’ end where they should have minimal effect on amplification. For both primers, a small proportion of sequences from the orders Gymnophiona and Caudata possessed mismatches within the last four bp of the 3’ end, so some small proportion of species may be missed from these orders. However, overall marker coverage for amphibian and turtle species was high.

**Figure 2.**
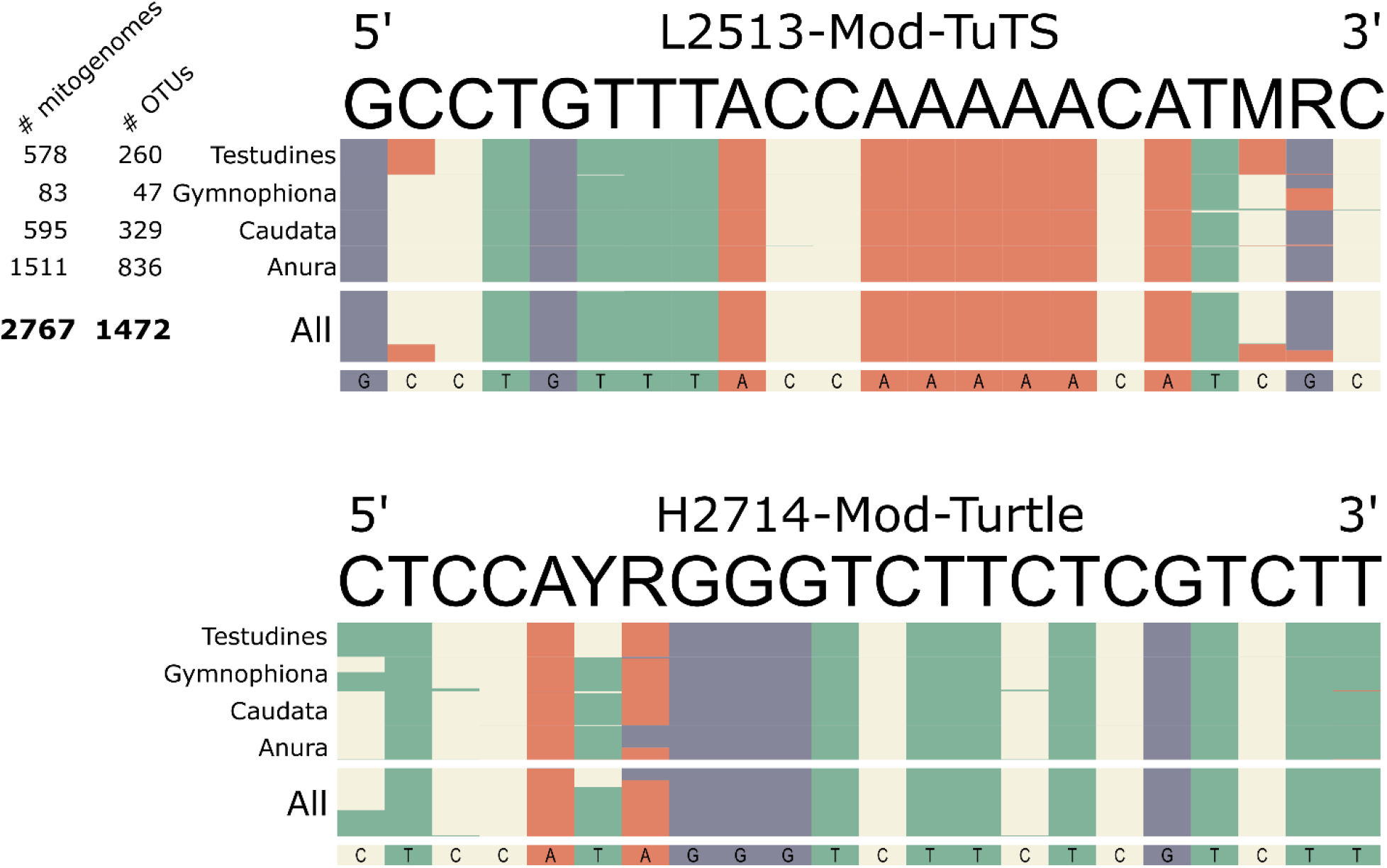
Plot of primers and primer region sequences from 2767 mitochondrial genomes from 9 fish orders. Columns represent the base pair composition at each position of the primer. # OTUs are the number of Operational Taxonomic Units after 97% clustering of mitochondrial genomes. Primer positions designated by R can match A or G, those designated by M can match A or C, and those designated by Y can match C or T.

For fish, we downloaded 8583 complete mitochondrial genomes from nine orders with PrimerMiner. These were clustered into 4762 OTUs representative of individual species. Nearly all fish sequences were a perfect match to both primers (Figure 3), with the exception of some mismatches near the 5’ end of the reverse primer H2714-Mod-Turtle which are unlikely to affect amplification. All sequences from the orders Siluriformes, Cypriniformes, and many from Characiformes possess a mismatch at the position of one of our degenerate bases 6 base pairs from the 5’ end of primer H2714-Mod-Turtle (Figure 3). It should be possible to replace the Y degenerate base from our primer with an H degenerate base to improve the efficiency of this primer for amplification of fish DNA for these orders if desired, but these mismatches are far enough from the 3’ to preclude amplification.

**Figure 3.**
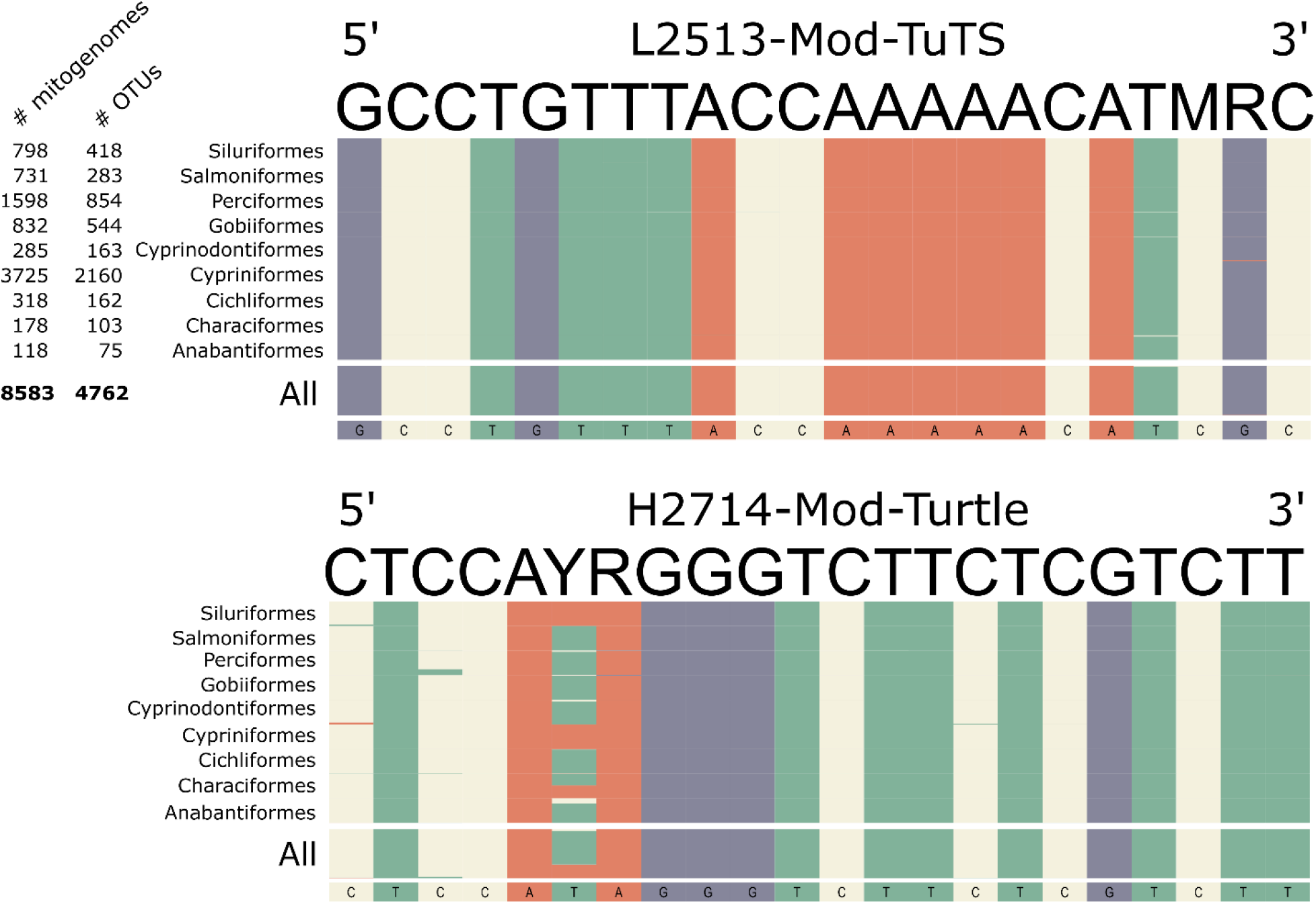
Plot of primers and primer region sequences from 8583 mitochondrial genomes from 9 fish orders. Columns represent the base pair composition at each position of the primer. # OTUs are the number of Operational Taxonomic Units after 97% clustering of mitochondrial genomes. Primer positions designated by R can match A or G, those designated by M can match A or C, and those designated by Y can match C or T.

## Discussion

We designed a novel primer set for amplification of amphibian, turtle and fish DNA for metabarcoding of environmental DNA samples. This marker performed well in detecting aquatic amphibian, turtle, and fish species from field-collected eDNA samples from California. Additionally, our *in-silico* validation suggests this marker should work well in detecting amphibian, turtle, and fish species globally.

Our novel metabarcoding marker generally performed better in detecting amphibian and turtle species from eDNA samples than an existing marker from the literature (*batra*; Valentini et al., 2016). This is most likely because our marker was designed specifically to perfectly match all amphibian and turtle species expected from the study area in California where eDNA samples were collected, while the published *batra* marker was designed for use globally. As a result, it is possible that the *batra* marker does not perfectly match all species from our California study area, resulting in reduced sensitivity for some species. The discrepancy was most notable for turtle species. The most commonly detected turtle species (*A. marmorata*) was detected in over 9-times more samples with our 16S-AmTu marker then with *batra*, and another turtle species (*T. scripta*) detected in 10 samples by 16S-AmTu was not detected at all by the *batra* marker. This discrepancy is not surprising, as our 16S-AmTu marker was specifically designed to amplify turtle DNA in addition to amphibians, whereas the *batra* marker was only designed for detection of species from the families Anura and Caudata. Thus, the *batra* maker clearly has lower sensitivity for detecting turtle species, even though it still provided some detections for two of the three species detected from eDNA samples with 16S-AmTu. Additionally, our 16S-AmTu marker detected 8/11 amphibian species from a greater number of samples than *batra*, and two amphibian species were only detected by the 16S-AmTu marker. However, three species were detected by eDNA samples more often by *batra* than by 16S-AmTu. One advantage of the *batra* marker is that it produces a much shorter amplicon than 16S-AmTu (52 bp vs. 196 bp), and this shorter amplicon length may confer greater sensitivity from degraded eDNA samples. Thus, it is possible that the *batra* marker could perform equally or better than our 16S-AmTu marker for detection of *Batrachia* species (but likely not turtles or fish) from eDNA in some situations, especially from locations other than central and southern California.

However, if the goal is detection of amphibians as well as turtle and fish species, our novel marker appears clearly preferable.

Despite being optimized for detection of amphibian and turtle species, the 16S-AmTu marker may work relatively well as a universal marker for vertebrate taxa in general. We demonstrate that this marker is also effective for detection of fish species, and other vertebrate taxa such as mammals were detected. As we were focused on aquatic species, we did not conduct a comprehensive evaluation of our primer sets for detecting terrestrial vertebrate species; however, it appears that this marker works well for detection of mammals as we detected 27 mammal species known from California from our eDNA samples (Table S4). Additionally, numerous OTUs belonging to bird taxa were detected; however, the taxonomic power of this marker for identifying bird taxa may be somewhat low, as some OTUs were a 100% match to reference sequences for bird species not residing in north America. The universality of our marker for detecting terrestrial vertebrates should be investigated further as some vertebrate groups are likely to be excluded, or the marker may not have high taxonomic power for some groups such as birds. For example, in a limited investigation into the ability of 16S-AmTu to detect reptile species other than turtles, the marker appeared to commonly have mismatches near the 3’ end of the reverse primer H2714-mod-turtle with many squamate species sequences. That said, OTUs belonging to two lizard species known from California, Southern alligator lizard (*Elgaria multicarinata*) and western fence lizard (*Sceloporus occidentalis*) were detected from eDNA samples. Additionally, a terminal mismatch was observed in all GenBank reference sequences for the order Crocodilia. Thus, it appears marker 16S-AmTu is unlikely to perform robustly for amplification of reptile DNA outside of the order Testudines.

In conclusion, the development and application of our novel metabarcoding marker 16S-AmTu represents an advancement in the field of (eDNA) research, particularly for aquatic vertebrate biodiversity assessment. Although metabarcoding markers exist for detecting amphibian, turtle, and fish species, we are not aware of any other markers designed to detect all three of these important aquatic vertebrate groups simultaneously. Running a single marker is highly preferable, and cost effective, compared to running multiple markers on the same samples. Our marker demonstrated robust performance in detecting amphibian, turtle, and fish species from field-collected eDNA samples from central California. Furthermore, in-silico evaluations suggest our novel marker has global applicability, enhancing its utility for broad-scale biodiversity monitoring. Notably, compared to an existing marker, our 16S-AmTu marker exhibited superior sensitivity in detecting target species, particularly turtles, underscoring its effectiveness for comprehensive aquatic vertebrate surveys. While our primary focus was on aquatic species, preliminary evidence suggests potential versatility for detecting terrestrial vertebrates, especially mammals, warranting further investigation into the marker’s applicability for terrestrial species. Overall, this study contributes a valuable tool for non-invasive monitoring of aquatic vertebrate communities, with implications for conservation, management, and ecological research.

## Acknowledgements

We would like to acknowledge the United States Department of Defense Strategic Environmental Research and Development Program (award # RC18-1034) for funding this work. We would like to thank field personnel involved in eDNA sample collection including Michael Andrews, Isidro Blanco, Jonathan Carmichael, Jessie Doyle, Kathleen Hicks, Gilbert Mak, Madison Ono, Savannah Pena, Andrew Reyna, Abigail Rivera, Megan Rodenbeck, Shaylea Stark, Emma Teall, Chris Terry, Kelsey Trammell, Jessica Turner, Fransico Villegas, Gretchen Wichman, Emily Wilkinson, Katharina Zimmermann. We thank Jacquelyn Hancock (Ft Hunter-Liggett), Rhys Evans (Vandenberg Space Force Base), Michael Moore and Michelle Wong (Camps Roberts and Camp San Luis Obispo), and Kristie Klose (Los Padres National Forest) for assisting with access and providing local knowledge. We also thank Lauren Scheinberg from the California Academy of Sciences and Carol Spencer from the Berkeley Museum of Vertebrate Zoology for providing access to tissue samples for sequencing.

