## Supplementary Information for "A novel degenerate primer set for eDNA metabarcoding of amphibians, turtles and fish"

**Table S1. Reference tissues sequenced for development of primers for metabarcoding.**

| **sample name** | **species name** | **common name** | **state** | **county** | **tissue source** | **GenBank accession number** |
| --- | --- | --- | --- | --- | --- | --- |
| RT173 | *Actinemys marmorata* | Western pond turtle | CA | Napa | [MVZ:Herp:164994](http://arctos.database.museum/guid/MVZ:Herp:164994) | MT135564 |
| RT174 | *Actinemys marmorata* | Western pond turtle | CA | Lake | [MVZ:Herp:164995](http://arctos.database.museum/guid/MVZ:Herp:164995) | MT135565 |
| RT108 | *Anaxyrus boreas halophilus* | Western toad | CA | Monterey | [CAS-206465](http://researcharchive.calacademy.org/research/herpetology/catalog/index.asp?xAction=getrec&close=true&CatalogNo=CAS+206465) | MT137267 |
| RT150 | *Anaxyrus boreas halophilus* | Western Toad | CA | Kern | [CAS-253026](http://researcharchive.calacademy.org/research/herpetology/catalog/index.asp?xAction=getrec&close=true&CatalogNo=CAS+253026) | MT137280 |
| RT97 | *Anaxyrus californicus* | Arroyo Toad | CA | San Diego | [CAS 175636](https://researcharchive.calacademy.org/research/herpetology/catalog/index.asp?xAction=getrec&close=true&CatalogNo=CAS+175636) | MT137260 |
| RT98 | *Anaxyrus californicus* | Arroyo Toad | CA | San Diego | [CAS 178994](https://researcharchive.calacademy.org/research/herpetology/catalog/index.asp?xAction=getrec&close=true&CatalogNo=CAS+178994) | MT137261 |
| RT99 | *Anaxyrus californicus* | Arroyo Toad | CA | San Diego | [CAS 179041](https://researcharchive.calacademy.org/research/herpetology/catalog/index.asp?xAction=getrec&close=true&CatalogNo=CAS+179041) | MT137262 |
| RT162 | *Anaxyrus californicus* | Arroyo Toad | Mexico | Baja Norte | [MVZ:Herp:145230](https://arctos.database.museum/guid/MVZ:Herp:145230) | MT135559 |
| RT112 | *Aneides lugubris* | Arboreal salamander | CA | Santa Cruz | [CAS-208660](http://researcharchive.calacademy.org/research/herpetology/catalog/index.asp?xAction=getrec&close=true&CatalogNo=CAS+208660) | MT137270 |
| RT168 | *Batrachoseps incognitus* | San Simeon slender salamander | CA | Monterey | [MVZ:Herp:251931](http://arctos.database.museum/guid/MVZ:Herp:251931) | MT135562 |
| RT145 | *Batrachoseps luciae* | Santa Lucia Mountains slender slamander | CA | Monterey | [CAS-252904](http://researcharchive.calacademy.org/research/herpetology/catalog/index.asp?xAction=getrec&close=true&CatalogNo=CAS+252904) | MT137279 |
| RT120 | *Batrachoseps nigriventris* | Black-bellied slender salamander | CA | San Luis Obispo | [CAS-214856](http://researcharchive.calacademy.org/research/herpetology/catalog/index.asp?xAction=getrec&close=true&CatalogNo=CAS+214856) | MT137272 |
| RT175 | *Chrysemys picta* | Painted turtle | NM | Sierra | [MVZ:Herp:250712](http://arctos.database.museum/guid/MVZ:Herp:250712) | MT135566 |
| RT176 | *Chrysemys picta* | Painted turtle | WA | Spokane | [MVZ:Herp:238581](http://arctos.database.museum/guid/MVZ:Herp:238581) | MT135567 |
| RT106 | *Ensatina eschscholtzii eschscholtzii* | Ensatina salamander | CA | Monterey | [CAS-205796](http://researcharchive.calacademy.org/research/herpetology/catalog/index.asp?xAction=getrec&close=true&CatalogNo=CAS+205796) | MT137266 |
| RT109 | *Ensatina eschscholtzii xanthoptica* | Yellow-eyed Ensatina salamander | CA | San Mateo | [CAS-207429](http://researcharchive.calacademy.org/research/herpetology/catalog/index.asp?xAction=getrec&close=true&CatalogNo=CAS+207429) | MT137268 |
| RT127 | *Lithobates catesbeiana* | America Bullfrog | CA | Fresno | [CAS-224776](http://researcharchive.calacademy.org/research/herpetology/catalog/index.asp?xAction=getrec&close=true&CatalogNo=CAS+224776) | MT137274 |
| RT151 | *Lithobates catesbeiana* | America Bullfrog | CA | Kern | [CAS-253042](http://researcharchive.calacademy.org/research/herpetology/catalog/index.asp?xAction=getrec&close=true&CatalogNo=CAS+253042) | MT137281 |
| RT155 | *Pseudacris cadaverina* | California chorus frog | CA | Los Angeles | [CAS-255413](http://researcharchive.calacademy.org/research/herpetology/catalog/index.asp?xAction=getrec&close=true&CatalogNo=CAS+255413) | MT137284 |
| RT165 | *Pseudacris cadaverina* | California chorus frog | Mexico | Baja Norte | [MVZ:Herp:145382](http://arctos.database.museum/guid/MVZ:Herp:145382) | MT135560 |
| RT238 | *Pseudacris hypochondriaca* | Baja California chorus frog | CA | Riverside | [CAS-200610](http://researcharchive.calacademy.org/research/herpetology/catalog/index.asp?xAction=getrec&close=true&CatalogNo=CAS+200610) | OP377739 |
| RT239 | *Pseudacris hypochondriaca* | Baja California chorus frog | CA | Riverside | [CAS-200616](http://researcharchive.calacademy.org/research/herpetology/catalog/index.asp?xAction=getrec&close=true&CatalogNo=CAS+200616) | OP377740 |
| RT111 | *Pseudacris sierra* | Sierran chorus frog | CA | San Luis Obispo | [CAS-208510](http://researcharchive.calacademy.org/research/herpetology/catalog/index.asp?xAction=getrec&close=true&CatalogNo=CAS+208510) | MT137269 |
| RT105 | *Rana boylii* | Foothill yellow-legged frog | CA | Santa Clara | [CAS-205752](http://researcharchive.calacademy.org/research/herpetology/catalog/index.asp?xAction=getrec&close=true&CatalogNo=CAS+205752) | MT137265 |
| RT131 | *Rana boylii* | Foothill yellow-legged frog | CA | Santa Clara | [CAS-226108](http://researcharchive.calacademy.org/research/herpetology/catalog/index.asp?xAction=getrec&close=true&CatalogNo=CAS+226108) | MT137275 |
| RT160 | *Rana boylii* | Foothill yellow-legged frog | CA | Amador | [CAS-259672](http://researcharchive.calacademy.org/research/herpetology/catalog/index.asp?xAction=getrec&close=true&CatalogNo=CAS+259672) | MT137285 |
| RT102 | *Rana draytonii* | California red-legged frog | CA | Contra Costa | [CAS-200290](http://researcharchive.calacademy.org/research/herpetology/catalog/index.asp?xAction=getrec&close=true&CatalogNo=CAS+200290) | MT137264 |
| RT125 | *Spea hammondii* | Western spadefoot | CA | Tulare | [CAS-223542](http://researcharchive.calacademy.org/research/herpetology/catalog/index.asp?xAction=getrec&close=true&CatalogNo=CAS+223542) | MT137273 |
| RT129 | *Spea hammondii* | Western spadefoot | CA | Stanislaus | [CAS-225300](http://researcharchive.calacademy.org/research/herpetology/catalog/index.asp?xAction=getrec&close=true&CatalogNo=CAS+225300) | OP131912 |
| RT166 | *Spea hammondii* | Western spadefoot | CA | San Joaquin | [MVZ:Herp:234174](http://arctos.database.museum/guid/MVZ:Herp:234174) | OP131913 |
| RT167 | *Spea hammondii* | Western spadefoot | CA | San Joaquin | [MVZ:Herp:234175](http://arctos.database.museum/guid/MVZ:Herp:234175) | MT135561 |
| RT203 | *Spea hammondii* | Western spadefoot | CA | San Luis Obispo | UCLA-132798 | OP115848 |
| RT204 | *Spea hammondii* | Western spadefoot | CA | San Luis Obispo | UCLA-132799 | OP115849 |
| RT213 | *Spea hammondii* | Western spadefoot | CA | Santa Barbara | UCLA-134538 | OP115857 |
| RT214 | *Spea hammondii* | Western spadefoot | CA | Santa Barbara | UCLA-134539 | OP115858 |
| RT218 | *Spea hammondii* | Western spadefoot | CA | Kern | UCLA-131797 | OP115862 |
| RT220 | *Spea hammondii* | Western spadefoot | CA | Orange | UCLA-132561 | OP115732 |
| RT224 | *Spea hammondii* | Western spadefoot | CA | Ventura | UCLA-131710 | OP115736 |
| RT225 | *Spea hammondii* | Western spadefoot | CA | Ventura | UCLA-131712 | OP115737 |
| RT226 | *Spea hammondii* | Western spadefoot | CA | San Diego | UCLA-131669 | OP115738 |
| RT227 | *Spea hammondii* | Western spadefoot | CA | San Diego | UCLA-131670 | OP115739 |
| RT228 | *Spea hammondii* | Western spadefoot | CA | Monterey | UCLA-133222 | OP115864 |
| RT229 | *Spea hammondii* | Western spadefoot | CA | Monterey | UCLA-133223 | OP115865 |
| RT200 | *Spea hammondii* | Western spadefoot | CA | Glenn | UCLA-9765 | OP115847 |
| RT118 | *Sternotherus odoratus* | Common musk turtle | FL | Taylor | [CAS-214343](http://researcharchive.calacademy.org/research/herpetology/catalog/index.asp?xAction=getrec&close=true&CatalogNo=CAS+214343) | MT137271 |
| RT154 | *Sternotherus odoratus* | Common musk turtle | GA | Macon | [CAS-255136](http://researcharchive.calacademy.org/research/herpetology/catalog/index.asp?xAction=getrec&close=true&CatalogNo=CAS+255136) | MT137283 |
| RT152 | *Taricha torosa* | California newt | GA | Kern | [CAS-253044](http://researcharchive.calacademy.org/research/herpetology/catalog/index.asp?xAction=getrec&close=true&CatalogNo=CAS+253044) | OP378122 |
| RT171 | *Taricha torosa* | California newt | GA | San Luis Obispo | [MVZ:Herp:236242](http://arctos.database.museum/guid/MVZ:Herp:236242) | OP378124 |
| RT159 | *Taricha torosa sierrae* | Sierra newt | GA | Madera | [CAS-259608](http://researcharchive.calacademy.org/research/herpetology/catalog/index.asp?xAction=getrec&close=true&CatalogNo=CAS+259608) | OP378123 |
| RT95 | *Taricha torosa sierrae* | Sierra newt | CA | Fresno | [CAS-208714](http://researcharchive.calacademy.org/research/herpetology/catalog/index.asp?xAction=getrec&close=true&CatalogNo=CAS+208714) | OP378125 |
| RT135 | *Trachemys scripta* | Pond slider | GA | Shasta | [CAS-227634](http://researcharchive.calacademy.org/research/herpetology/catalog/index.asp?xAction=getrec&close=true&CatalogNo=CAS+227634) | MT137276 |
| RT177 | *Trachemys scripta elegans* | Red-eared slider | NM | Sierra | [MVZ:Herp:265667](http://arctos.database.museum/guid/MVZ:Herp:265667) | MT135568 |
| RT142 | *Xenopus laevis* | African clawed frog | CA | San Francisco | [CAS-244034](http://researcharchive.calacademy.org/research/herpetology/catalog/index.asp?xAction=getrec&close=true&CatalogNo=CAS+244034) | MT137278 |
| RT153 | *Xenopus laevis* | African clawed frog | CA | San Francisco | [CAS-253186](http://researcharchive.calacademy.org/research/herpetology/catalog/index.asp?xAction=getrec&close=true&CatalogNo=CAS+253186) | MT137282 |

**Table S2. Tissue DNA samples included in pools for testing of metabarcoding marker.**

| **Sample Name** | **species** | **source** |
| --- | --- | --- |
| RT85 | *Ambystoma californiense* | [CAS-211677](https://researcharchive.calacademy.org/research/herpetology/catalog/index.asp?xAction=getrec&close=true&CatalogNo=CAS+211677) |
| RT97 | *Anaxyrus californicus* | [CAS 175636](https://researcharchive.calacademy.org/research/herpetology/catalog/index.asp?xAction=getrec&close=true&CatalogNo=CAS+175636) |
| RT162 | *Anaxyrus californicus* | [MVZ:Herp:145230](https://arctos.database.museum/guid/MVZ:Herp:145230) |
| RT168 | *Batrachoseps incognitus* | [MVZ:Herp:251931](http://arctos.database.museum/guid/MVZ:Herp:251931) |
| RT145 | *Batrachoseps luciae* | [CAS-252904](http://researcharchive.calacademy.org/research/herpetology/catalog/index.asp?xAction=getrec&close=true&CatalogNo=CAS+252904) |
| RT120 | *Batrachoseps nigriventris* | [CAS-214856](http://researcharchive.calacademy.org/research/herpetology/catalog/index.asp?xAction=getrec&close=true&CatalogNo=CAS+214856) |
| RT108 | *Anaxyrus boreas halophilus* | [CAS-206465](http://researcharchive.calacademy.org/research/herpetology/catalog/index.asp?xAction=getrec&close=true&CatalogNo=CAS+206465) |
| RT175 | *Chrysemys picta* | [MVZ:Herp:250712](http://arctos.database.museum/guid/MVZ:Herp:250712) |
| RT173 | *Emys marmorata* | [MVZ:Herp:164994](http://arctos.database.museum/guid/MVZ:Herp:164994) |
| RT174 | *Emys marmorata* | [MVZ:Herp:164995](http://arctos.database.museum/guid/MVZ:Herp:164995) |
| RT148 | *Ensatina eschscholtzii* | [CAS-252913](https://researcharchive.calacademy.org/research/herpetology/catalog/index.asp?xAction=getrec&close=true&CatalogNo=CAS+252913) |
| RT117 | *Ensatina eschscholtzii croceator* | [CAS-213213](https://researcharchive.calacademy.org/research/herpetology/catalog/index.asp?xAction=getrec&close=true&CatalogNo=CAS+213213) |
| RT106 | *Ensatina eschscholtzii eschscholtzii* | [CAS-205796](http://researcharchive.calacademy.org/research/herpetology/catalog/index.asp?xAction=getrec&close=true&CatalogNo=CAS+205796) |
| RT103 | *Ensatina eschscholtzii xanthoptica* | [CAS-203543](https://researcharchive.calacademy.org/research/herpetology/catalog/index.asp?xAction=getrec&close=true&CatalogNo=CAS+203543) |
| RT111 | *Pseudacris sierra* | [CAS-208510](http://researcharchive.calacademy.org/research/herpetology/catalog/index.asp?xAction=getrec&close=true&CatalogNo=CAS+208510) |
| RT155 | *Pseudacris cadaverina* | [CAS-255413](http://researcharchive.calacademy.org/research/herpetology/catalog/index.asp?xAction=getrec&close=true&CatalogNo=CAS+255413) |
| RT100 | *Rana draytonii* | [CAS-200288](http://researcharchive.calacademy.org/research/herpetology/catalog/index.asp?xAction=getrec&close=true&CatalogNo=CAS+200288) |
| RT157 | *Rana draytonii* | [CAS-257329](https://researcharchive.calacademy.org/research/herpetology/catalog/index.asp?xAction=getrec&close=true&CatalogNo=CAS+257329) |
| RT105 | *Rana boylii* | [CAS-205752](http://researcharchive.calacademy.org/research/herpetology/catalog/index.asp?xAction=getrec&close=true&CatalogNo=CAS+205752) |
| RT160 | *Rana boylii* | [CAS-259672](http://researcharchive.calacademy.org/research/herpetology/catalog/index.asp?xAction=getrec&close=true&CatalogNo=CAS+259672) |
| RT127 | *Rana catesbeiana* | [CAS-224776](http://researcharchive.calacademy.org/research/herpetology/catalog/index.asp?xAction=getrec&close=true&CatalogNo=CAS+224776) |
| RT125 | *Spea hammondii* | [CAS-223542](http://researcharchive.calacademy.org/research/herpetology/catalog/index.asp?xAction=getrec&close=true&CatalogNo=CAS+223542) |
| RT166 | *Spea hammondii* | [MVZ:Herp:234174](http://arctos.database.museum/guid/MVZ:Herp:234174) |
| RT118 | *Sternotherus odoratus* | [CAS-214343](http://researcharchive.calacademy.org/research/herpetology/catalog/index.asp?xAction=getrec&close=true&CatalogNo=CAS+214343) |
| RT161 | *Taricha torosa* | [CAS-259679](https://researcharchive.calacademy.org/research/herpetology/catalog/index.asp?xAction=getrec&close=true&CatalogNo=CAS+259679) |
| RT95 | *Taricha torosa sierrae* | [CAS-208714](http://researcharchive.calacademy.org/research/herpetology/catalog/index.asp?xAction=getrec&close=true&CatalogNo=CAS+208714) |
| RT135 | *Trachemys scripta* | [CAS-227634](http://researcharchive.calacademy.org/research/herpetology/catalog/index.asp?xAction=getrec&close=true&CatalogNo=CAS+227634) |
| RT177 | *Trachemys scripta elegans* | [MVZ:Herp:265667](http://arctos.database.museum/guid/MVZ:Herp:265667) |
| RT142 | *Xenopus laevis* | [CAS-244034](http://researcharchive.calacademy.org/research/herpetology/catalog/index.asp?xAction=getrec&close=true&CatalogNo=CAS+244034) |

**Table S3. Tissue DNA samples from “exotic” species not known from North America used for positive controls in eDNA metabarcoding libraries.**

| **Sample Name** | **species name** | **common name** | **country of origin** | **source** |
| --- | --- | --- | --- | --- |
| RT179 | Sclerophrys regularis | African common toad | Republic of Cameroon | [CAS-253847](https://researcharchive.calacademy.org/research/herpetology/catalog/index.asp?xAction=getrec&close=true&CatalogNo=CAS+253847) |
| RT180 | Lissemys punctata | Indian flapshell turtle | Pakistan | [CAS-232082](https://researcharchive.calacademy.org/research/herpetology/catalog/index.asp?xAction=getrec&close=true&CatalogNo=CAS+232082) |
| RT181 | Pelomedusa gehafie | Eritrean helmeted turtle | Eritrea | [CAS-262594](https://researcharchive.calacademy.org/research/herpetology/catalog/index.asp?xAction=getrec&close=true&CatalogNo=CAS+262594) |
| RT182 | Ptychadena sp. | grassland frog | Eritrea | [CAS-262432](https://researcharchive.calacademy.org/research/herpetology/catalog/index.asp?xAction=getrec&close=true&CatalogNo=CAS+262432) |
| RT183 | Tomopterna kachowskii | Kachowski's Sand Frog | Eritrea | [CAS-262451](https://researcharchive.calacademy.org/research/herpetology/catalog/index.asp?xAction=getrec&close=true&CatalogNo=CAS+262451) |
| RT184 | Tylototriton shanorum | newt | Myanmar | [CAS-230933](https://researcharchive.calacademy.org/research/herpetology/catalog/index.asp?xAction=getrec&close=true&CatalogNo=CAS+230933) |

**Table S4. Mammal species detected from environmental DNA samples. Taxonomic identification was based on top hit from NCBI BLAST matching a mammal species known from California.**

| **Species name** | **Common name** |
| --- | --- |
| Canis latrans | coyote |
| Castor canadensis | American beaver |
| Cervus canadensis nannodes | Tule elk |
| Chaetodipus californicus | California pocket mouse |
| Didelphis virginiana | common opossum |
| Felis catus | domestic cat |
| Sciurus griseus | western gray squirrel |
| Ictidomys tridecemlineatus | thirteen-lined ground squirrel |
| Lynx rufus | bobcat |
| Microtus californicus | California vole |
| Neotoma fuscipes | dusky-footed woodrat |
| Neotoma lepida | desert woodrat |
| Neotoma macrotis | big-eared woodrat |
| Odocoileus virginianus | white-tailed deer |
| Ondatra zibethicus | muskrat |
| Onychomys sp. | grasshoper mouse |
| Peromyscus boylii | brush mouse |
| Peromyscus crinitus | canyon mouse |
| Peromyscus maniculatus | deer mouse |
| Procyon lotor | raccoon |
| Puma concolor | puma |
| Rattus rattus | black rat |
| Sus scrofa | domistic pig |
| Sylvilagus bachmani | brush rabbit |
| Tamias spp. | chipmunk |
| Urocyon cinereoargenteus | grey fox |
| Ursus americanus | black bear |
